# Codon based co-occurrence network motifs in human mitochondria

**DOI:** 10.1101/092262

**Authors:** Pramod Shinde, Camellia Sarkar, Sarika Jalan

## Abstract

Human mitochondrial DNA (mtDNA) generates a highly divergent pool of alleles even within species that have dispersed and expanded in size. On top of that codon position bias plays a relative role in prompting population wide sequence based variations. Codon positions are immensely studied as independent evolutionary genetic markers, however co-evolution of codon positions are still largely unexplored. Using 18,411 genome samples, we analyze nucleotide co-occurrences over mitochondrial DNA of human sub-populations covering most of the mtDNA diversity on the earth. Our codon based network motif investigation reveals that codon position is one of the critical factor to form higher order motifs. It is interesting to report a different evolution of Asian genomes than those of the rest which is divulged by motifs in non-coding regions. Most notably, we demonstrate that there are preferences of codon position co-evolution with respect to codon position sensitivity implying the selective role of evolutionary processes on fundamental formation of genetic code in terms of codon bias evolution. These network based findings not only recapitulates the established facts regarding the importance of each codon position in providing codon bias but also highlight their preferential role in resulting correlated mutations. Moreover, codon based nucleotide co-occurrence provides framework to investigate important evolutionary insights into human mitochondrial evolutionary genomics.

## 1 Introduction

“Nature’s stern discipline enjoins mutual help at least as often as warfare. The fittest may also be the gentlest.” as proclaimed by Theodosius Dobzhansky on mankind evolution, like galaxy formations, evolution of human remain the generation long grit of thinkers. Non-recombining loci, such as the maternally inherited mitochondrial DNA have been known to provide a richer estimation insight into ancestral human genetic variations during evolution^1^, as mtDNA usually pass intact from generation to generation and hence they preserve a simpler record of their history^2^, ^3^. Independent dispersal of evolutionary signatures in mtDNA and their extent of conservation in human sub-populations has been delivering a fertile research area of human evolution since last two decades^4^, ^5^. Further, the genetic code is degenerate. Each amino acid is encoded by particular set of codons; the choice of codons within a genome is not random but are evolutionary selected to incorporate codon bias^6^, ^7^. Role of codon positions are immensely studied as independent evolutionary genetic markers^8^, ^9^, however co-evolution of codon positions are still largely unexplored. Moreover, genetic markers must be interacting with one another, at least in the loose sense that both influence the same phenotype^10^. In essence, the evolutionary behavior of the genome often involves co-operative changes in the variable genome positions^11^ and results in various evolutionary genomic discernments which we precisely captured in this Letter using network model.

Complex network science revolves around the hypothesis that the behavior of complex systems can be elucidated in terms of the structural and functional relationships between their constituents by means of a graph representation^12^. The network framework provides cue into whether the structural environment confers opportunities for or constraints on individual node action^13^. Network motifs, considered to be topologically distinct interaction patterns, are known to be fundamental feature of networks and represent the simplest building blocks^14,15^. In biological networks, they represent meaning to protein family as well as to pathway conservation^16^ which has been shown theoretically and experimentally as well^17^. Furthermore despite the tremendous advancements in the field of network theory, very few have taken nucleotide based genomic co-occurrences into consideration^18,19^ whereas codon based genomic co-occurrences have not been described previously to the best our knowledge. Co-occurrences of codon and non-codon positions between the genomes across sub-populations is expected to help in gaining insights into co-changes in genomes as well as conservation of these co-changes across sub-populations since these co-changes are said to be largely dispositioned during evolutionary events such as climatic changes, migration events, genetic traits etc^20^.

Here, we consider around 18,000 human mitochondrial genomes which covers most of the mtDNA diversity of human population on the earth. Our analysis show that variable nucleotide positions among mitochondrial genomes residing in different human sub-populations implement the independent mtDNA evolution with respect to its geographical dispensation. Construction of nucleotide co-occurrence networks with respect to different human sub-populations provides fundamental insights into genetic code in terms of codon bias evolution which we study in depth using analysis of network motifs. Network motifs based on codon positions give the evolutionary signature of association between codon positions present within and between human sub-populations. Revealing importance of each codon position in mitochondrial evolution, our codon based network motif analysis yields evolutionary preferences of codon positions in the formations of lower and higher order motifs. Most interestingly, we find that most of the lower order motifs are formed within Asian genomes indicating a different evolutionary mechanism.

## 2 Results

### 2.1 Statistics of Variable sites

#### 2.1.1 Variome is well conserved and independently maintained

Variable site is a sequence variation occurring at a particular genomic position where a single nucleotide (A, T, G or C) in a genome differs between members of a species (Fig. 1a). There are total 17,174 variable sites observed in all the genome groups which make total 7065 unique variable sites out of 16,881 bps genome size. It means that 42% nucleotide positions of human mtDNA are incidentally atleast once mutated till date. We find that each genome group has different sizes of variome (V), also the large portion of these variomes in each genome group occurs at similar genome positions than that of another genome group(s) (Table 1 and Fig. 2b). It reiterates the fact that mutational sites might have descended from co-evolutionary mechanisms among genomes of different genome groups. Further, we find that the substantial portion of variomes (21%) is common between all genomes groups (Fig. 2b). It suggests that the majority of variable sites of these geographical variomes are commonly incidented across human population. Further, we look for how variomes are shared between the set of four, three and two genome groups. We find that the largest portion of variomes are shared among {*AS,AM,AF,EU*}, {*AF,AS,EU*} and {*AM,AS,AF*} genome groups (Fig. 2b), indicating that more number of variable sites are co-occurred between human population habitating in AF, AM, AS and EU geographical regions than that of with OC geographical region. It also suggests that AS and EU genome groups share the highest number of variable sites. In sum, count of common as well as different variable sites which are found between different genome groups implicate that different evolutionary mechanisms might be functional among genomes of different genome groups. Variable sites introspect more robust functionalities to the mitochondrial genome^21^ and above observations show importance of variable sites to shape not only genotypic polymorphism but also the associated phenotypic variation since variable sites in variomes are differently incidented among genomes of different population samples. Here, we also perceive that mitochondrial genomes have more sequence diversions residing among some genome groups than others. Nevertheless, observations with variable sites reiterate the fact that molecular sequence based diversions in terms of single nucleotide polymorphism are very well conserved across genome groups as well as independently maintained within genome groups.

**Figure 1.**
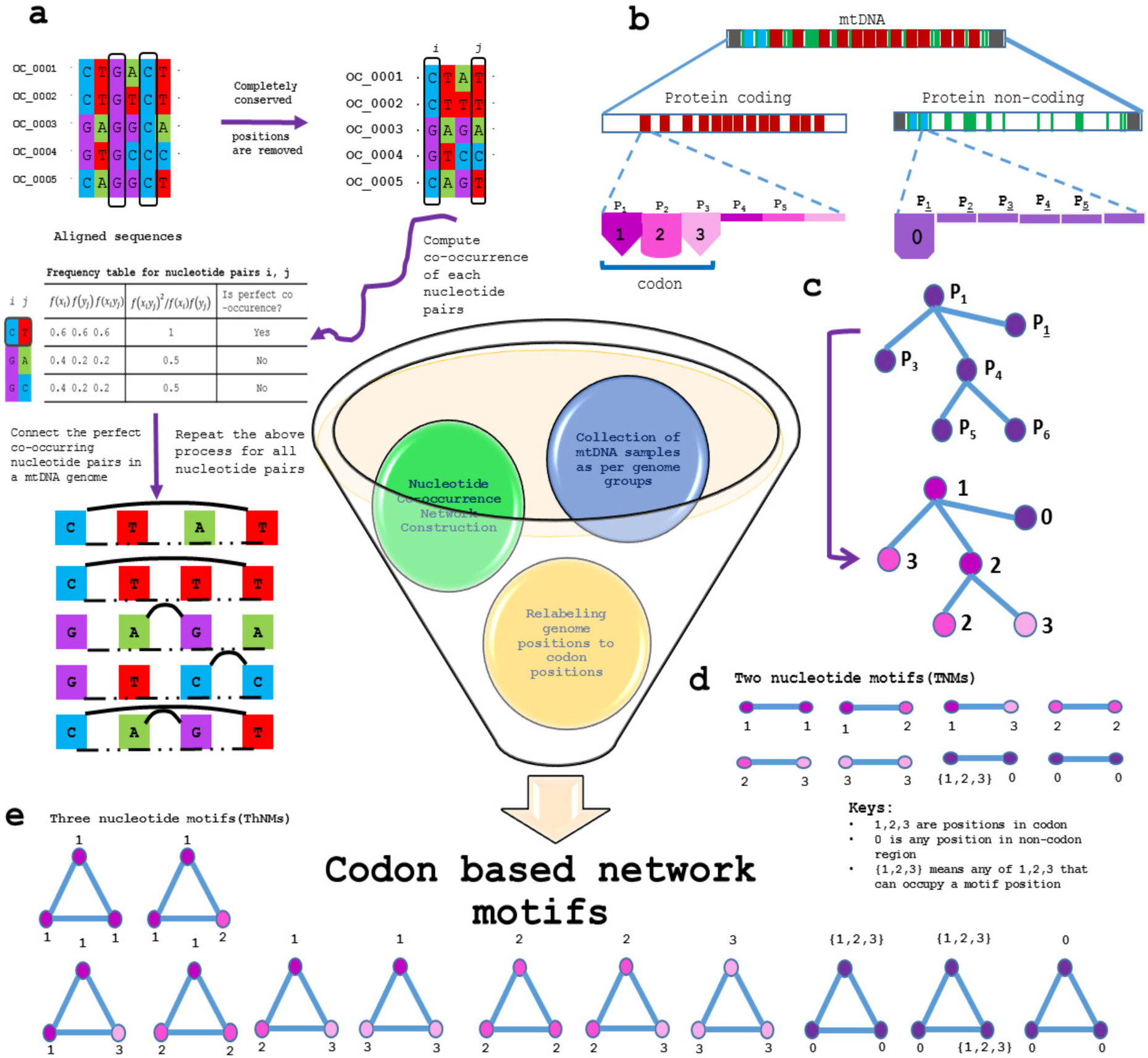
An overview of codon based nucleotide co-occurrence network analysis. (a) Flow diagram depicts the schematic process of nucleotide co-occurrence network construction. Stepwise, it involves the removal of conserved nucleotide positions and the selection of variable sites, followed by preparation of frequency table for each pairing and then selection of perfect co-occurring nucleotide pairs^18^. As a result, nucleotide co-occurrence network for each genome is constructed (Supplementary Fig. S1), likewise there are 18,411 networks considered for the study. (b) The whole genome of mitochondria consists of protein coding and protein non-coding regions. Translation of protein coding gene begins at translation start site and further elongation of protein synthesis continues with reading a set of three nucleotides termed as codon. Codon positions are identified with respect to corresponding nucleotide positions. (c) Genome positions are relabeled as the codon positions for each nucleotide co-occurrence network. For instance, *P*_1_ is present in protein coding region which is relabeled as 1 whereas *P*<u>_1_</u> is present in protein non-coding region which is relabeled as 0. Likewise, entire network is relabeled for further analysis. (d) and (e) display different types of two and three nucleotide motifs possible in nucleotide co-occurrence networks. Circle represents node and numbers nearby the circle represent the position of codon. We define these motifs structures as homogeneous and heterogeneous motif structures. Homogeneous motifs are the structures where all motif positions are occupied by the same codon position e.g., 1–1, 2–2, 3–3 as well as 1–1-1, 2–2-2, 3–3-3. Heterogeneous motifs are formed when motif positions are occupied by different codon positions e.g., 1–2 as well as 1–2-3.

**Table 1.** Data and network statistics. *S* represents sample size or number of genomes, *NV* represents number of variable sites or size of variome. 〈*NCo*〉 and 〈*E*〉 represent average number of nodes and average number of edges across networks of a genome group, respectively where as 〈*k*〉 represents average degree of the networks of a genome group.

| Genome Group | $S$ | $N_V$ | $\langle N_{Co}(\pm) \rangle$ | $\langle E(\pm) \rangle$ | $\langle k \rangle$ |
| --- | --- | --- | --- | --- | --- |
| Africa (AF) | 2323 | 3255 | 914 (5) | 967 (6) | 2 |
| America (AM) | 1692 | 2990 | 967 (4) | 1211 (5) | 2 |
| Asia (AS) | 5715 | 4898 | 426 (5) | 474 (19) | 2 |
| Europe (EU) | 7142 | 4466 | 455 (2) | 473 (1) | 2 |
| Oceania (OC) | 1539 | 1564 | 470 (5) | 334 (8) | 2 |

**Figure 2.**
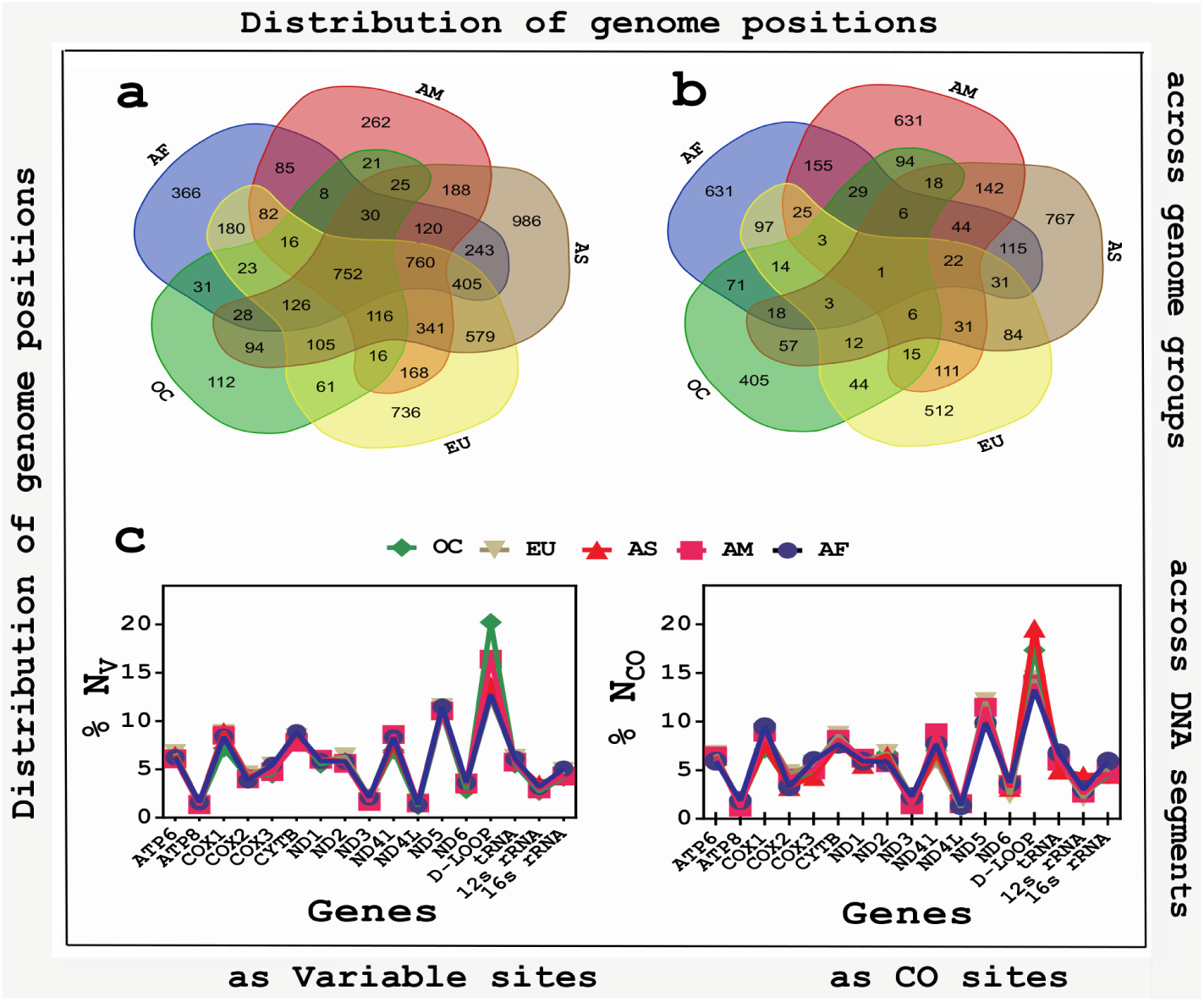
Distribution of genome positions as variable sites and CO sites across genome groups and DNA segments. (**a** and **b**) Variable sites as well as CO sites tend to occur at similar genome positions in different set of genome groups such as two, three, four and all five. There are certain variable sites which only occur at particular genome group. (**c**) Percentage of V (left panel) and CO (right panel) occurred in both protein coding and non-coding DNA segments of mitochondrial genome. There are total 22 tRNA genes present in human mitochondria, we term all tRNA segments as one tRNA to find their collective role. Count of both V and CO show similar patterns of increase and decrease.

#### 2.1.2 Co-occurring variable sites show region specific variations

We investigate co-occurrence of variable sites within each genome since these sites are thought to be co-evolved^18^. When we say variable sites are co-occurred (*C_f_* = 1), it means that the presence of a particular nucleotide at a genome position is associated with a particular nucleotide at another genome position. It is to note that a small portion of variable nucleotide sites participate as perfect co-occurring variable sites (CO) which constitute nodes of the network (Table 1). Unlike variable sites, CO do not share more number of sites in common between genome groups (Fig. 2a and b). Extending these observations, we note that distribution of V and CO show similar patterns in increase and decrease with respect to their occurrence among individual DNA segments as well as among all genome groups (Fig. 2c). It implicates that whether variable sites are participating as co-occurring variable sites or not, they are conserved. Further, we find that V and CO occur at both protein coding and non-coding regions (Fig. 2c). These results not only suggest that single nucleotide polymorphism can occur throughout mtDNA but also highlight that these population genomics signatures are widely conserved among all DNA segments across genome groups. Regional variations accumulate mtDNA diversity purely attributed to genetic drift in each genome group^22,23^ and above observations suggest that region specific variations persist in resulting co-occurrence of sequence based variations.

#### 2.1.3 CO pairs display intra- and inter- DNA adaptation

Analysis of CO pairs provides an essential understanding of relationship between two genome locations (Fig. 3). Genetic adaptation in response to selection on polygenic phenotypes may occur via subtle allele frequency shifts at many loci or they are resultant of effector mutations at one or many places^25^. CO pairs can be formulated within a particular mitochondrial gene (intra-gene) or between two mitochondrial genes (inter-gene). We enumerate CO pairs and calculate the number of CO pairs formed across the length of mtDNA. There are less number of CO pairs formed between intra-gene segments than inter-gene segment (Fig. 3a). It shows the importance of much anticipated polygenic co-evolution among mtDNA genes. The relationship between correlated mutations and spatial proximity has not only been found between residues in the same protein but also between residues in different proteins^26,27^. Along these lines, we count for the CO pairs formed between each intra- and inter-DNA segments (Fig. 3a). D-Loop has maximum CO pairs between intra-gene segments as well as it forms the least number of CO pairs with rest of the genome. COX1 forms the highest number of CO pairs occurring between inter- DNA segments. We also find that tRNA regions form much lesser (0.7%) intra-gene CO pairs, but at the same time they form CO pairs with the almost all polypeptide genes. It suggests that evolution of tRNA genes is largely controlled by recipient *i.e*., polypeptide genes.

**Figure 3.**
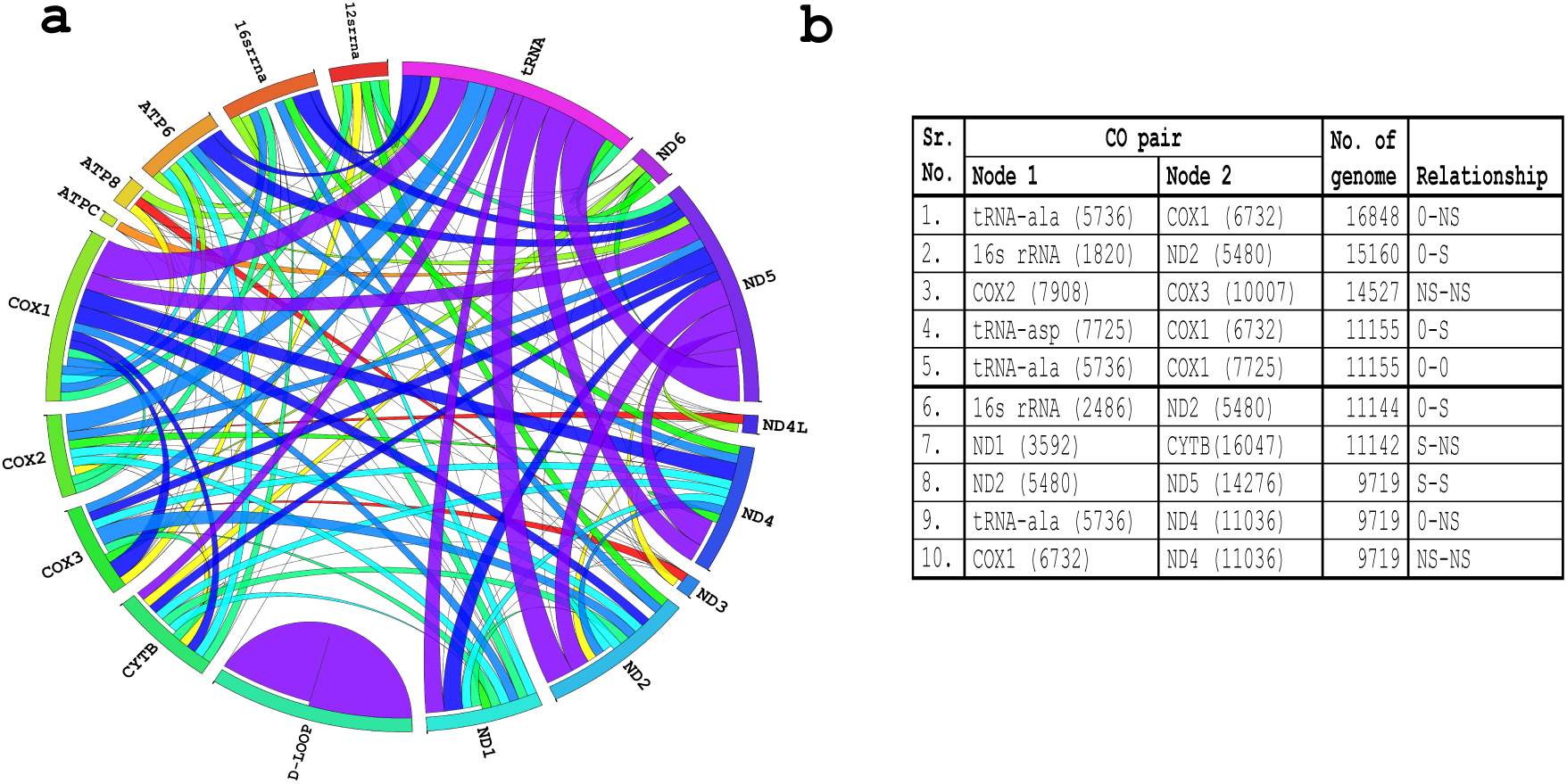
Statistics of CO pairs. (**a**) Circos plot illustrates shared CO pairs between polypeptide gene, non-coding, rRNA and tRNA regions. Link between two DNA segments represent these two DNA segments share CO pairs and the width of connection represent the amount of CO pairs. For instance, D-Loop has maximum number intra DNA segment CO pairs. (**b**) Table of predominant CO pairs existing among all genomes showing CO pairs occur between genome positions *i.e.*, node 1 and 2 and hence represents association between two genome positions (shown in associated braces). Further, we label them with codon positions which represents the relation between different types of mutations *i.e.*, synonymous (S) and non-synonymous (NS) mutations. Mutations among non-codon regions are also termed as S mutations^24^, but to make it distinguished from codon position 3 based mutation, we label mutations at genome sites at non-codon regions as 0. These associations between predominant CO pairs are of S-NS, S-S, NS-NS types between polypeptide genes as well as of S-0 and NS-0 between polypeptide gene and non-gene DNA segments.

Apart from tRNA regions, we find that polypeptide genes viz. ND5, COX1 also form CO pairs with almost all genes (Fig. 3a). At the same time, ND1 and COX1 form the least number of intra-gene CO pairs. All this suggest that there can be associations between nucleotide positions of single or multiple genes and individual DNA segment has particular preference to form CO pairs within as well as outside the DNA segment. These associations can be corollary of different environmental effects such as climatic and geography based events resulted into natural selection occurring in particular gene^28^. Both gene wide association studies as well as molecular structure based interaction studies have broadly shed light on the fact^21^ and we hope that perfect nucleotide co-occurrence may provide insights into intra- and inter- gene adaptation.

Moreover, it would be interesting to look for functional insights into how individual CO pair is formulated. In order to explore on this, we note down CO pairs which are predominant amongst all CO pairs as well as present in atleast two genome groups and picked up the top 10 pairs to understand their biological insights (Fig. 3b). These CO pairs are formed between inter-DNA segments and exist between polypeptide genes (e.g., COX2-COX3, ND1-CYTB, ND2-ND5, COX1-ND4) as well as polypeptide gene and non-gene regions (e.g., COX1-tRNA, ND4-tRNA etc). These pairs are evolutionary more conserved since they co-occur in maximum number of genomes across genome groups and are thought to provide genotypic insights in terms of what mutation they encode for. On the basis of relation between codon positions as well as type of mutations displayed by predominant CO pairs (Fig. 3b), it is possible to say that there can be any mutational preference to form CO pair between gene and non-gene regions as well as between different codon pairs (see legends of Fig. 3b). But it is interesting to observe that most of the CO pairs, present in atleast two genome groups, are formed either between polypeptide genes or between tRNA and polypeptide gene, implementing selective role of coding DNA to form CO pairs (illustrated in Supplementary Information).

### 2.2 Co-occurrence networks exhibit same average connectivity

We constructed nucleotide co-occurrence network for each genome, where all variable sites forming CO pairs constitute the nodes and the edges represent co-occurring nucleotide positions. Network size (*N*) is found to be nearly same within each genome group (Table 1) which is expected since these genomes have high sequence similarities as much as 99% and they differ only at variable genome sites. Though genomes between genome groups also have high sequence similarity, still *N* is found to be different which is intuitive as there are different *N_V_* present in each genome group (Table 1) which can be considered as one of the inherent properties of that genome group. Further, it is surprising to observe that average connectivity (〈*k*〉) of each co-occurrence network which represents the average number of co-varying partners per nucleotide position is found to be same for all the networks across all genome groups and attends a small value *i.e.*, 〈*k*〉 ***=*** *2* (Table 1). All this implicates that though co-evolution is an economic evolutionary mechanism^29^, still co-evolution of nucleotide positions is widely conserved across all mitochondrial co-occurrence networks. Further, we study how different codon positions are associated with identification of network motifs for 13 mtDNA polypeptide genes *i.e.*, codon regions and in like manner we extend our understanding by discerning network motifs between codon regions and other parts of the genome *i.e.*, non-codon regions.

### 2.3 Codon positions make network motifs

Network motifs are complete sub-graphs of order two or more. The codon positions in each motif share common biological functions. It means that particular codon position will have same sensitivity across the length of genome. For instance, codon position 1 will have high sensitivity irrespective of codon placement at any location in genome. Typically, every codon position is controlled by a particular type of mutational forces working at that genome position. In general cases, the resultant is termed as ‘single nucleotide polymorphism’, most of the variable codon positions in a motif are regulated precisely with the S and NS type of mutation in codon regions^30^. For instance, an example of a motif is the set of alterations in amino acid due to change in codon within binding site of protein regulated by NS mutations upon selection force^30,31^. We find that a number of motifs as large as in the real nucleotide co-occurrence network occurs very rarely in that of corresponding randomized networks (Fig. 4 and 5). The significant finding is that region specific interactions among motifs exist and that they seem to partition the group of codon positions into biologically meaningful clusters which we study precisely in subsequent.

**Figure 4.**
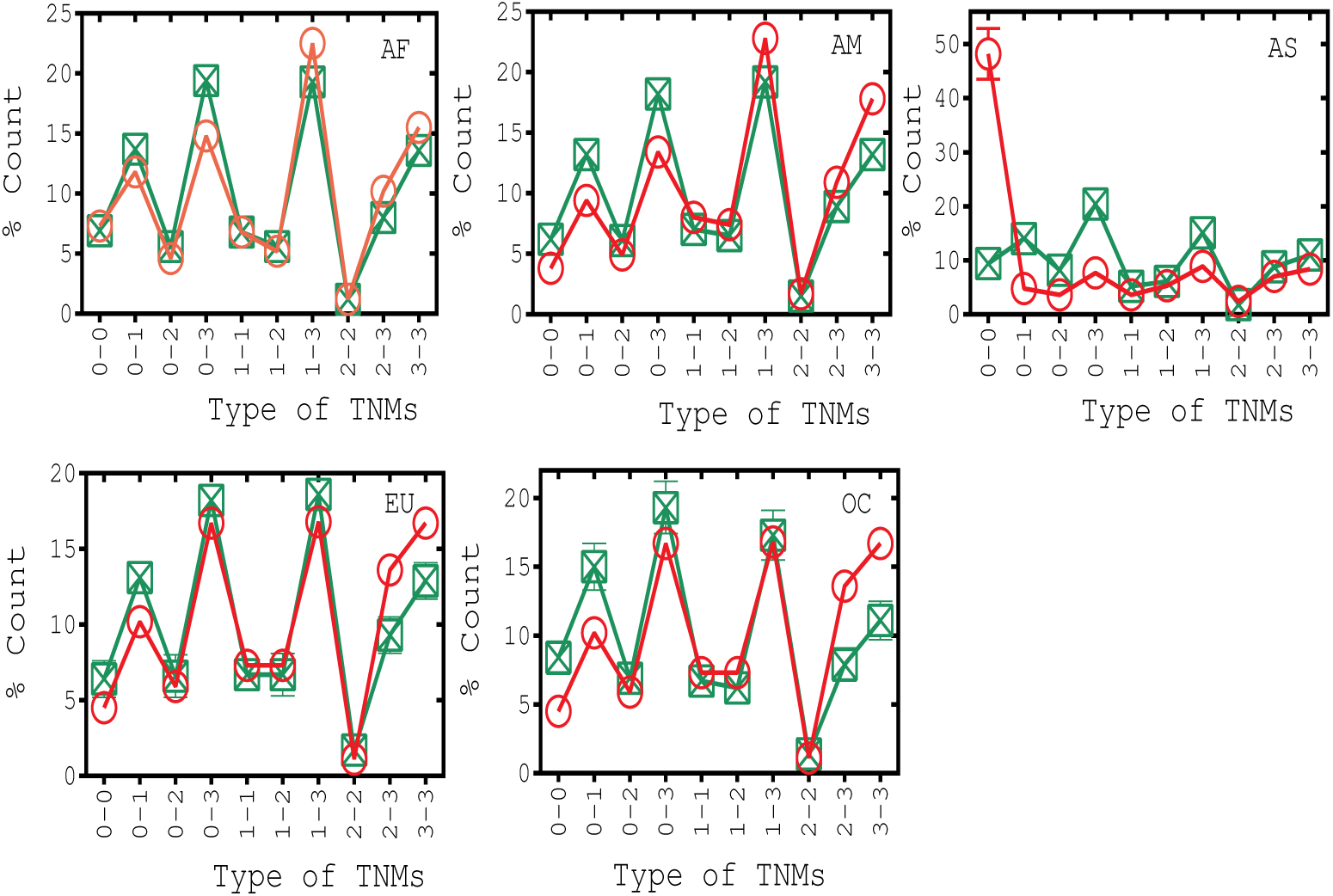
Distribution of TNMs in different genome groups. These motifs are made up of different codon positions such as 1, 2 and 3 whereas 0 in motif represents the nucleotide position of non-codon region. There are 10 possible TNM connected patterns in which 0–0 is found among non-codon regions, 0–1, 0–2 and 0–3 are found between codon and non-codon region whereas remaining TNMs are found among codon regions. Circles and squares denote motif count in nucleotide co-occurrence and randomized networks, respectively. We take 10,000 realizations of randomized networks for each genome group.

**Figure 5.**
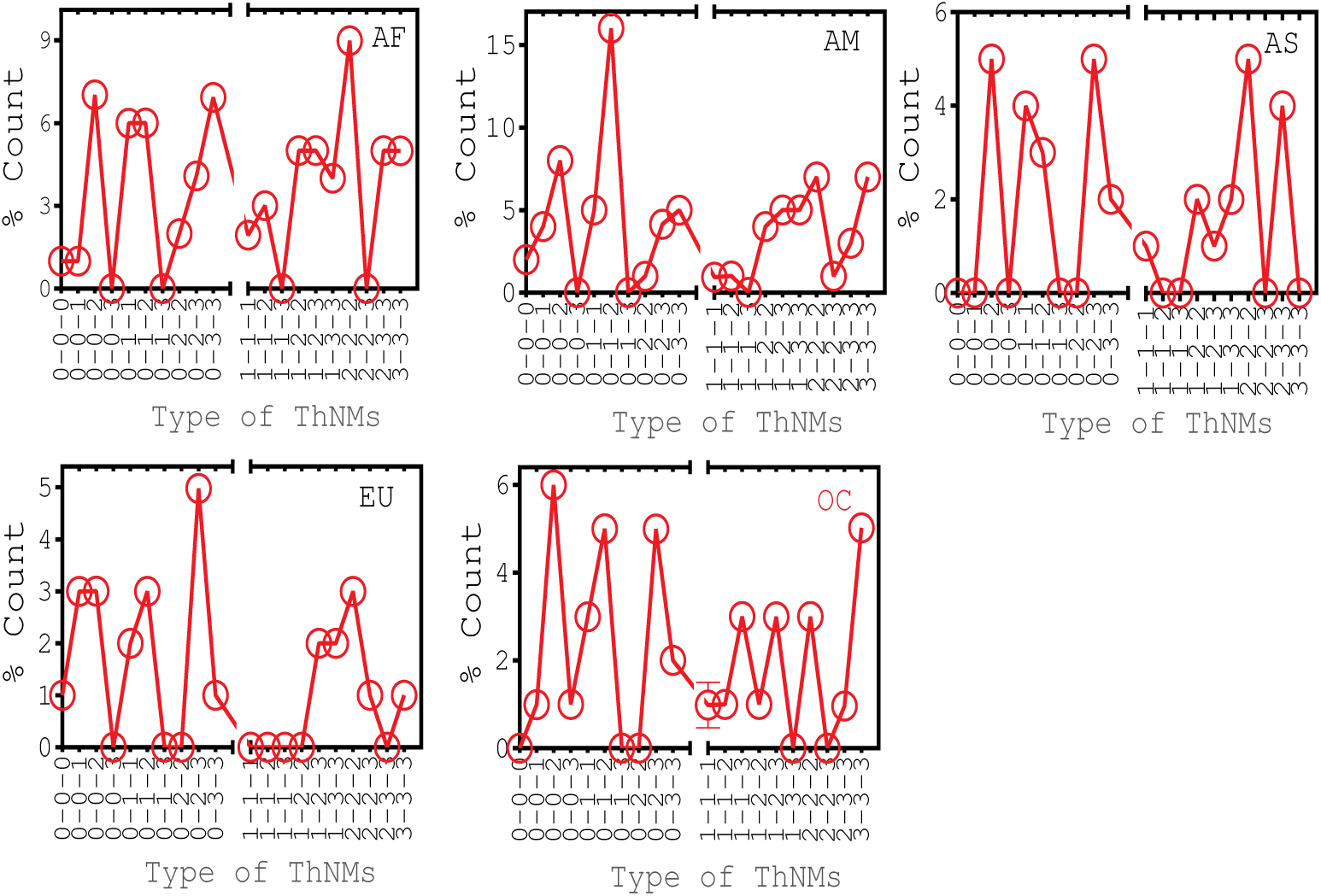
Distribution of ThNMs in different genome groups. There are 20 possible ThNMs. We term codon positions 1, 2 and 3 as least-, mid- and most- sensitive codon positions, respectively. These motifs are made up of different codon positions such as 1, 2 and 3 whereas 0 in motif represents the nucleotide position of non-codon region. Circles denote motif count in nucleotide co-occurrence networks. Randomized networks show significantly lesser number of ThNMs (Supplementary Fig. S2).

#### 2.3.1 Two nucleotide motifs (TNMs)

The fact that the network motifs among codon region appear at frequencies much higher than expected at random suggests that they may have specific functions and formation of TNMs is not merely random. This observation also indicates an ascendancy of codon positions in the motif formation. It is interesting to observe that most of the TNMs are formed where codon position 3 is acting as one of the motif nodes (Fig. 4), overtoning the puissance of codon position 3 in the motif formation. It is also observed that codon position 3 motifs are different in count than those of the corresponding randomized networks (Fig. 4). It implicates that there might be an evolutionary preference given to the motifs with codon position 3 in the codon regions. These results provide an essential understanding regarding synonymous and nonsynonymous mutations which is the nerve of the codon position bias^32^. It urges that less sensitive codon sites co-evolve higher and result in more S-S associations (3–3). Alternatively, this observation is also supported by lesser NS-S (1–3 and 2–3) as well as NS-NS (1–1, 1–2 and 2–2) associations than S-S associations. It not only reiterates the fact that nonsynonymous mutations are found in lesser number than the synonymous mutations^33^ but also suggests that most-sensitive codon sites exhibit lesser preference to co-evolve among themselves as compared to those of the least-sensitive codon sites in short range conservations.

Interestingly, non-codon region has lesser motifs than those of the randomized networks for all genome groups except AS (Fig. 4). 0–0 motifs in AS genome group are found to be significantly higher than that of the randomized networks. In line particularly, we analyze TNMs present among non-codon regions *i.e.*, tRNAs, rRNAs and D-Loop regions (Table 2). Surprisingly most of 0–0 motifs in AS genome group are formed within intra D-Loop region. It implicates a different evolution arising in the Asian mitochondrial genomes. Also, there are more number of TNMs present among intra- individual D-Loop (D-Loop: D-Loop) and intra- RNA (e.g., tRNA: rRNA) regions than inter- D-Loop and RNA regions (e.g., D-Loop: tRNA) in all genome groups (Table 2). D-Loop and rRNA are known to be the signatures of population genomics variations in most of the mtDNA associated studies^34^ and similar fact might be observable in the associations among non-codon positions as well since count of motifs in non-codon positions is found to be different than that of the random counterpart. Most provocatively, analysis of TNMs implicates the dominance of codon position 3 in human mitochondrial genomes as well as an exceptional role of D-Loop in Asian genomes.

**Table 2.** Percentage pairwise associations between 0–0 motif pairs among non-codon regions. DNA segment pair is a set of two non-codon mitochondrial DNA segments which consists of co-occurring variable sites each one from either DNA segment. For instance, DNA segments rRNA and tRNA have 32% of variable sites shared among each other in AF geographical region.

| DNA segment pair | AF(%) | AM(%) | AS(%) | EU(%) | OC(%) |
| --- | --- | --- | --- | --- | --- |
| rRNA : tRNA | 32 | 38 | 0 | 25 | 40 |
| rRNA : D-Loop | 0 | 0 | 0 | 0 | 4 |
| tRNA : D-Loop | 0 | 0 | 0 | 0 | 0 |
| rRNA : rRNA | 18 | 21 | 3 | 42 | 12 |
| tRNA : tRNA | 18 | 13 | 1 | 8 | 20 |
| D-Loop : D-Loop | 32 | 28 | 96 | 24 | 24 |

#### 2.3.2 Three nucleotide motifs (ThNMs)

We detect locally dense regions in the network which are thought to be evolutionarily conserved^35^. There have been several different approaches to identify clustering based network motifs and these studies have provided distinct outcomes with respect to the resulting network motifs^36^. The main conclusion, however, is that nucleotide co-occurrence networks tend to be locally connected^18,19^ whereas network motifs are entirely separable from the rest of the network. In fact, we find that many identified these complete sub-graphs are nested within each other.

ThNMs in genome networks are found to be significantly higher than that of the randomized networks which is not surprising. However, the interesting result is that most of the ThNMs are formed when atleast one node of the motif belongs to the codon region (Fig. 5). It reveals the importance of codon positions in long range conservations. It is well known that evolution of mitochondrial genome involves optimum use of each nucleotide position especially codon position for functional as well as evolutionary adaptations^37^. Above results suggest that functional associations between protein coding genes are evolutionarily more conserved than associations between other parts of mtDNA. In addition, it would be intriguing to study how these codon positions co-evolve amongst as well as with other codon positions to devise long range conservations. We find that predominant ThNMs are homogeneous triplets of codon positions 2 and 3 except for AS genomes where 3–3-3 is negligible. It is interesting to observe that mid-sensitive codon position associations (2–2-2) form predominant homogeneous triplet though codon position 2 forms lesser number of TNMs and ThNMs with other codon positions (Fig. 4 and 5). It indicates that mid-sensitive codon positions favor co-association with themselves while forming higher order motifs. Further as we discussed earlier, the least-sensitive codon positions tend to form TNMs amongst themselves and it would be interesting to study how these codon positions form higher order motifs. Particularly, we analyze the association of motif 3–3 with codon positions 1 and 2, and it has resulted in no specific preference to form triplets with any codon position. In a way, it implicates that though short range conservation of least-sensitive codon positions are in abundance, they are limited to form long range conservations with other codon positions. Further, we find that homogeneous triplet with codon position 1 is formed in lesser number as compared to other homogeneous motifs. This observation is consistent with the observation of 1–1 homogeneous TNMs. It implicates that codon position 1 involves less in homogeneous motif formation. This is a vital observation since codon position 1 is a highly sensitive site for mutation and hence it is known to be highly conserved^6^. It impels that the most-sensitive genome sites such as codon position 1 are less preferred to co-evolve among themselves.

Moreover, we analyze ThNMs involving non-codon positions and find that the count of 0–0-0 ThNMs is less which is consistent with the lesser count of 0–0 motifs (except in AS). It implicates that there is a lesser preference to form motifs in non-codon regions. Also, it is observed that cumulatively more number of motifs are formed when codon positions are associated with non-codon positions. It incriminates the preference of codon position to get co-evolved with non-codon region of mtDNA. Besides that it is interesting to observe that codon position 2 is co-associated with 0–0 motifs with as much as 12 ± 4% of total ThNMs where as at the same time 0–0 motifs show less co-association preference with codon position 1 and 3. In addition, we study heterogeneous triplets between codon and non-codon regions and find that 0–1-2, 0–2-3 types are the prominent heterogeneous triplets (Fig. 4). It implicates that though 1–3 is the predominant order two motif, it shows a lesser preference to associate with non-codon positions. Further, it would be interesting to study how these heterogeneous TNMs are associated with codon positions. We find that both 1–2 and 1–3 motifs have more preference to form a higher order motifs with codon positions 2 and 3 than that of the codon position 1. Similarly, we find that motif 2–3 shows preference to form higher order motifs with codon positions 1 and 3 than that of the codon position 2. To summarize, these results suggest that there might be a particular preference assigned to each heterogeneous pair to form higher order motifs.

## 3 Discussion

Because mitochondrial metabolism is highly sensitive to environmental conditions, particularly temperature variations, one can speculate that the human mtDNA evolved in response to environmental or body temperature to maintain optimal function of mitochondria^38^. This fact is notably observed in our analysis of variable sites and motif where they show heterogeneity across genome groups. Conceding coding DNA is driver of genome evolution^6^, codon based motif analysis demonstrates different co-occurrence preferences among codon positions in genome evolution. The nucleotide co-occurrence network is made up of variable sites which are known to be the signatures of population variations^1^. We find that these sites are differently distributed among genome groups implementing the independent evolution of genomes among human sub-populations^39^. Overall observations with V and CO are attributed into causality factors such as natural selection processes which might have played a role in shaping human regional mtDNA variation^23^. Relabeling these variable sites with the corresponding codon positions provide an efficient way to study genomic co-evolution which are reported to be resultant of codon position bias^40^. Codon based motif analysis by subdividing network into several connected patterns reveals a more comprehensive view about the correspondence between genomic co-changes and characteristics of the evolution among mtDNA in human sub-population. Mitochondrial genome is the most suitable evolutionary model to study codon based nucleotide co-occurrence since it has only one non-codon DNA region whereas other RNA genes are directly known to co-evolve with mtDNA protein coding genes^1^, ^2^.

Lesser count of TNMs among non-codon positions as compared to that of the random counterpart suggests that evolutionary conservation prefers to happen at codon regions which have direct role in controlling phenotype. Interestingly, 0–0 motifs are significant in AS region resulting in unique feature shown by co-occurrence network motifs. Particularly, we find that these 0–0 motifs are resultant of intra D-Loop and intra RNA co-evolution which have been extensively reported as evolutionary signatures of human population genomics^34^. Moreover, comparison with the corresponding randomized networks suggests that higher order motif formation is not merely random and is one of the properties of nucleotide co-occurrence. Adding to this, there are least number of ThNMs formed within non-codon regions whereas it is possible to form ThNMs only when there is a codon position involved. Our analysis reveals that (1) the codon position is one of the essential ingredient to form higher order motifs indicating that long range nucleotide conservations across mtDNA are consequence of phenotypic associations *i.e.*, resultant of protein coding DNA. (2) Least-sensitive codon positions have a higher mutational ability as they are more involved as variable sites. They also form large number of TNMs implementing that synonymous mutations might have consequence to get evolved with codon as well as non-codon positions. Though codon position 3 delivers silent mutations, it is still reported to provide genetic signature(s) to population genomics studies. (3) Mid-sensitive codon positions display the specificity to form long range evolutionary conservations that too with homogeneous structure formation and with non-codon region. Also, most-sensitive sites prefer less to co-evolve and vice versa. These results alt the particular act of evolutionary mechanisms on each codon position to get differentially associated with other part of genome. In all, above results depict how codon position co-occurrences participate in codon bias evolution which supports the fact that codon bias is functionally demanding and provide basis to genetic code formation^8^.

Essentially, our TNMs based analysis suggest that although each codon position has significant role in network motif formation and these structures are evolutionarily adapted among genome groups, still the overall statistics of motif structures is remained to be conserved for lower order motif formation throughout genome groups. There are other principal genetic mechanisms that contribute to the evolution of new functions to mtDNA such as functional divergence of mtDNA genes, functional interactions between mtDNA and nuclear DNA etc^41^. No doubt many of these facts have contributed in co-changes among mtDNA but also are reported to be further resulted into stabilizing mutations among other part of mtDNA which provide an independence to the mtDNA evolution^41^. Other than that strong mutualism amongst genes of mtDNA provides an independent evolutionary functionality to mtDNA^2, 42^. Owing to these facts, our predictions are more relevant in relating admixture of codon and non-codon associations and provide a much needed codon glance to genomic positions, a novel network approach to understand genome evolution. Further, it would be interesting to extend the observations of the current study in comparison of human mtDNA with those of primates and other mammals^43^. Our approach can be readily generalized to any type of nucleotide co-occurrence network to improve on evolutionary understanding of codons and their associations across prokaryotic as well as eukaryotic genes.

## 4 Methods

### 4.1 Genome sequences

We retrieve all genome sequences in fasta file format from the Human Mitochondrial Database (Hmtdb)^44^ which provides a comprehensive, integral, non-redundant set of mtDNA genomes from geographically diverse human populations (http://www.hmtdb.uniba.it/hmdb/; data collected on Feb 16, 2016). The ascertainment set comprises 18,411 genome sequences from the five world continents, including 2323 African, 1692 American, 5715 Asian, 7142 European and 1539 Ocean genomes (see Supplementary Information). We termed each of them as a genome group.

### 4.2 Construction of mitochondrial nucleotide co-occurrence networks

For each of these five sub-population genome datasets, we construct the nucleotide co-occurrence network in which nodes represent nucleotides at specific positions in the genome sequence and edges between nodes represent co-occurring nucleotide pairs (Fig. 1) as following. (1) Since, the current study is based on the analysis of specific nucleotide position in the genome sequence, we consider sequence data which is already end to end aligned, maintaining each nucleotide position in genome sequence uniform. (2) All conserved nucleotide positions within samples of a genome group are removed, we are thus left behind with the variable nucleotide positions. We term a set of variable sites in a genome group as variome. The count of variable sites (*N_V_*) is given in Table 1. (3) Using variable nucleotide positions, we first calculate the frequency of occurrence of all the nucleotide pairs *f*(*x_i_y_j_*) = *N*(*x_i_y_j_*)/*M* where, *N*(*x_i_y_j_*) denotes the number of co-occurrence pair (*x_i_y_j_*) at position (*i, j*) and *M* denotes total number of samples in a genome group. Second, we calculate the frequency of individual occurrence of single nucleotides *f*(*x_i_*) = *N*(*x_i_*)/*M* and *f*(*y_j_*) = *N*(*y_j_*)/*M* where, *N*(*x_i_*) and *N*(*y_j_*) denote the number of single nucleotides at their respective positions i and j.^18^ (4) Co-occurrence of two nucleotides (*CO*) at position (*i, j*) is denoted as,

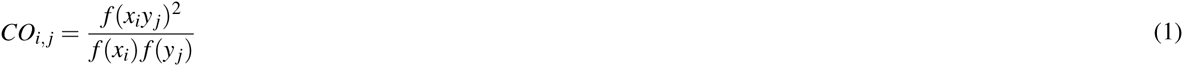

For perfect co-occurring variable sites, we define the adjacency matrix of the corresponding network as,

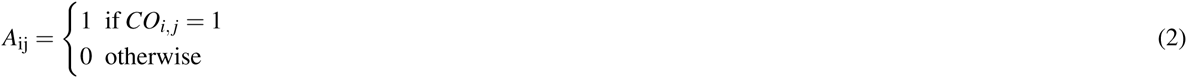

As each genome sequence has its own information of co-occurring nucleotide pairs, there are total *M* networks generated for each genome group.

### 4.3 Network motifs based on codon position

All genome files were parsed to extract 13 polypeptide genes, 22 tRNAs, two rRNAs and non-coding segments by aligning and comparing them with the standard reference sequences *i.e.*, the revised Cambridge Reference Sequence (rCRS, GenBank id: NC012920.1). We define network motifs based on the assignment of codon position to the genome position. As it is described in schematic Fig. 1b, the group of three nucleotide position in a gene region makes a codon. After construction of the nucleotide co-occurrence network, we re-label nodes with the position of codon. We refer rCRS in order to perform relabeling of codon positions. It is to note that HmtDb has also used the same reference sequence to perform the multiple sequence alignment of all the human mitochondrial genome which we consider for the current study. Further, we identify codon position of each genome position with respect to the reading frame of individual gene using their sequence position alignment with rCRS. In this way, each nucleotide position in protein coding gene regions (13 polypeptide genes) are relabeled with either 1, 2 or 3 and the rest of the genome positions *i.e.*, non-codon positions are labeled as 0. We enumerate two nucleotide (TNMs) and three nucleotide (ThNMs) network motifs for each nucleotide co-occurrence network (see legends of Fig. 1b). After detailing all the motifs based on their positions in codons in different networks of each genome group, we perform various statistical analysis of TNMs and ThNMs.

### 4.4 Generation of randomized networks

For a stringent comparison to randomized networks, we generated random networks with precise information of number of nodes and number of connections that of corresponding real nucleotide co-occurrence network. Construction of randomized networks and their comparison with corresponding real networks allow to estimate the probability that a randomized network with certain constraints has of belonging to a particular architecture, and thus assess the relative importance of different network architecture. This information is collected by taking an average of number nodes (N) and average degree (〈*K*〉) of networks present in each geographical group. The randomized network of size N and 〈*K*〉 is constructed using the Erdős Renyi random network model by connecting each pair of nodes with the probability (p)^45^.

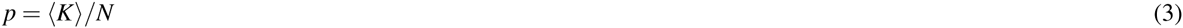

In this way, 10,000 randomized networks are constructed for each genome group. We analyze TNMs and ThNMs for each real nucleotide co-occurrence network and compare them with those of the corresponding randomized networks^12^.

## 5 Acknowledgements

SJ. is grateful to Department of Science and Technology (DST), Government of India grant EMR/2014/000368 for financial support. We appreciate interesting discussions with Prof. Michael Hiller, MPI-GCI, Germany and Dr. Obul Reddy Bandapalli, German Cancer Research Insititute, Germany. P.S. acknowledges DST for the INSPIRE fellowship (IF150200) and Dr. Analabha Basu, NIBMG, India for initial discussions during NGBT 2015, Hyderabad, India. Authors thank Complex Systems Lab members for timely help and fruitful discussions.

## Contributions

S J. and PS. conceived and designed the project. PS. and C.S. generated the data and performed all the analysis. S J. supervised the study. All the authors wrote and approved the manuscript.

